# Opposites attract: Multiple evidence of sexual antagonistic coevolution driving extreme male-biased sexual size dimorphism in *Panopeus meridionalis*

**DOI:** 10.1101/2024.04.03.588019

**Authors:** N.E. Farias, P. Ribeiro, J.P. Lancia, T. Luppi

## Abstract

Explanations for the evolution of male-biased size dimorphism (MBSSD) traditionally focus on male competition and female choice, overlooking the alternative that larger males may be better at coercing females into mating. While displaying considerable diversity, ‘true crabs’ (Eubrachyura) share common traits that theoretically should promote the evolution of coercive mating strategies. Despite this, there is a conspicuous lack of studies investigating this aspect. We investigated several reproductive and life history traits of *Panopeus meridionalis* (a mud crab that exhibits extreme MBSSD) to assess whether the specific set of characters is consistent with the hypothesis of sexual antagonistic coevolution at place. We found that the high MBSSD is likely driven by sexual conflict, where males use their size to coerce females into mating. Experimental matings involved male aggression towards females. Females first resist male attempts, but are ultimately subdued. Mating is relatively brief and there is no evident pre or post copulatory guarding. The female reproductive tract lacks complex structures for long-term sperm storage or manipulation, and given the small size of seminal receptacles related to male sperm load capacity, it is unlikely for females to store sperm from multiple partners. All considered, the evidence suggests that females have limited control over paternity and support the existence of an intrinsically coercive mating system in *P. meridionalis*. We propose this species as an interesting model for studying the resolution of sexual conflict through antagonistic coevolution and selection in the highly diverse group of true crabs.

## INTRODUCTION

The role of sexual selection as the primary determinant of sexual differences in traits is a topic of debate and may vary depending on the taxa and methods used for its study (Janicke & Fromonteil, 2021). Character displacement between the sexes resulting from natural selection acting differentially in each sex, which is often reffered as Sexually Antagonistic Selection (Parker, 2006; Connallon & Hall, 2018; De Lisle, 2019), should not be overlooked as either the sole or a concurrent force driving sexually divergent evolution. Indeed, trait variation between sexes is very often shaped by a synergistic combination of drivers of natural and sexual selection, each bearing variable weight along their co-evolutionary trajectories (Wilhelm *et al*., 2015; De Lisle, 2019). A prime example of this is sexual size dimorphism (Connallon & Hall, 2018; Janicke & Fromonteil, 2021).

In systems with male-biased sexual size dimorphism (MBSSD), it’s often observed that larger males secure a disproportionately high number of matings. Traditionally, this was attributed to either female choice or superior performance of larger males in competing for mates, both components of the sexual selection. Sexually antagonistic selection is less frequently invoked, but there is substantial published evidence of its importance in driving SSD (Cox & Calsbeek, 2009; Connallon & Clark, 2014). Alternatively, a third compelling explanation often overlooked, is that larger males possess enhanced abilities to subdue and coercively inseminate females, thereby introducing an element of sexual conflict (Shine & Mason, 2005). When reproductive interests of males and females collide, the conditions are present for the development of sex-specific morphological and behavioral traits aimed at maximizing their individual reproductive success, even at the expense of the other sex (Arnqvist & Rowe, 2005). For example, in the many taxa where males invest considerably less in their offspring compared to females, they are likely to achieve substantial fitness gains by coercing females into mating (Trivers, 1972; Parker, 1979). This sexual conflict is expected to drive the evolution of male traits that enhance their coercive abilities, while simultaneously fostering those female traits that promote resistance or avoidance of forced mating or any other alternative that mitigate potential negative fitness consequences, such as post-copulatory cryptic choice (Burke & Holwell, 2021) or ecological displacement. If this dynamic persists, sexes may engage in rapid divergent evolution of the involved traits or the coevolution between males and females that is driven by sexual conflict) leading to extreme sexual dimorphisms (Parker, 2006). Lastly, the degree of the resulting differences in male and female traits, however, is constrained by the limitations of the inherent genetic linkage between the sexes, ensuring that the reproductive system as a whole (including mating strategies and behaviors) remains viable.

Distinguishing between actual male coercion and cryptic female choice or cooperation often poses a considerable challenge and requires experimental manipulation in most cases. Strictly, mating is considered coercive if it decreases female fitness (Clutton-Brock & Parker, 1995), something extremely hard to prove in natural conditions (De Lisle, 2019). A less strict definition, that sexual coercion occurs whenever there is ‘…(male) action biasing mating in a way that subverts an existing (female) mating preference…’ (Snow & Prum, 2023), might be useful for distinguishing whether a given trait evolved in a scenario of sexual coercion. Even then, a conclusion may require the integration of multiple sources of evidence to assess whether a particular set of traits is consistent with what would be expected for an explicitly coercive mating system at play (Eberhard, 2002).

Decapod crustaceans became model organisms for the study of the evolution of sexual systems (Duffy & Thiel, 2007). The Eubrachyura or ‘true crabs’, the most speciose group of decapods, exhibit complex life history traits, elaborate reproductive strategies, and some peculiar behaviors in mate attraction and mating itself (McLay and Becker, 2015). In true crabs, males tend to be larger than females, fertilization is internal and females bear the whole costs of parental care, as they brood the eggs until hatch without any aid from the male (Hines, 1992; Rodgers *et al*., 2011). During copulation, the male deposits ejaculates, consisting of spermatophores and seminal fluids, directly into the female seminal receptacles, a multipurpose organ that serves for insemination, sperm storage, fertilization, and oviposition. This structure endows females with the potential to exert control over the reproductive outcome (González-Gurriarán et al., 1998; Rorandelli et al., 2008; McLay & López Greco, 2011) by providing the anatomical stage for postcopulatory selection by cryptic female choice and sperm competition. All the above considered and leaving ecological drivers aside, there are good theoretical grounds to anticipate male sexual harassment and coercive mating to be a favored male strategy to overcome female reluctance to mate. However, there is a notable scarcity of studies investigating sexual coercion as an evolutionary driving force of importance for this taxon.

This study focuses on *Panopeus meridionalis*, a mud crab species that exhibits marked male-biased sexual size dimorphism. We hypothesize that this extreme MBSSD is linked to a reproductive system featuring overt coercive mating. We investigate various aspects of male and female biology, anticipating a set of traits that should reflect a balance between intersexual differences driven by sexual antagonistic coevolution and the genetic linkage complementarity. Ultimately, we synthesize the biological information herein generated and scrutinize its coherence with the hypothesis of the existence of an overt coercive mating operating in the species.

## MATERIALS AND METHODS

### Collection site and methods

The collection of individuals primarily took place between September and March of 2012 and 2013 in Mar Chiquita, Argentina (37° 35’S, 57° 26’W, Fig. 1). Mar Chiquita is a brackish coastal lagoon of roughly 46 km^2^ with low amplitude tides not exceeding 1 m (Spivak et al., 1994). Despite anecdotal records in the region, a stable population of *P. meridionalis*, featuring regularly observed ovigerous females within the lagoon, has not been reported until 2005 (Spivak & Luppi, 2005). This habitat is a saltmarsh dominated by the sea grass *Spartina* spp. and heavily influenced by the activity of the burrowing crab, *Neohelice granulata*. At the lagoon’s inlet, a series of small to medium-sized orthoquartzite rock jetties has been built (Fig. 1). Some of these boulders are scattered across the lower intertidal and upper subtidal zones, with the latter becoming visible only during exceptionally low tides or when strong winds push the water towards the sea. Although *P. meridionalis* is rare in Mar Chiquita, it is readily found beneath the rocks in this boulder-strewn area. All individuals utilized in this study were collected by hand during normal low tides from this specific location.

**Fig 1.**
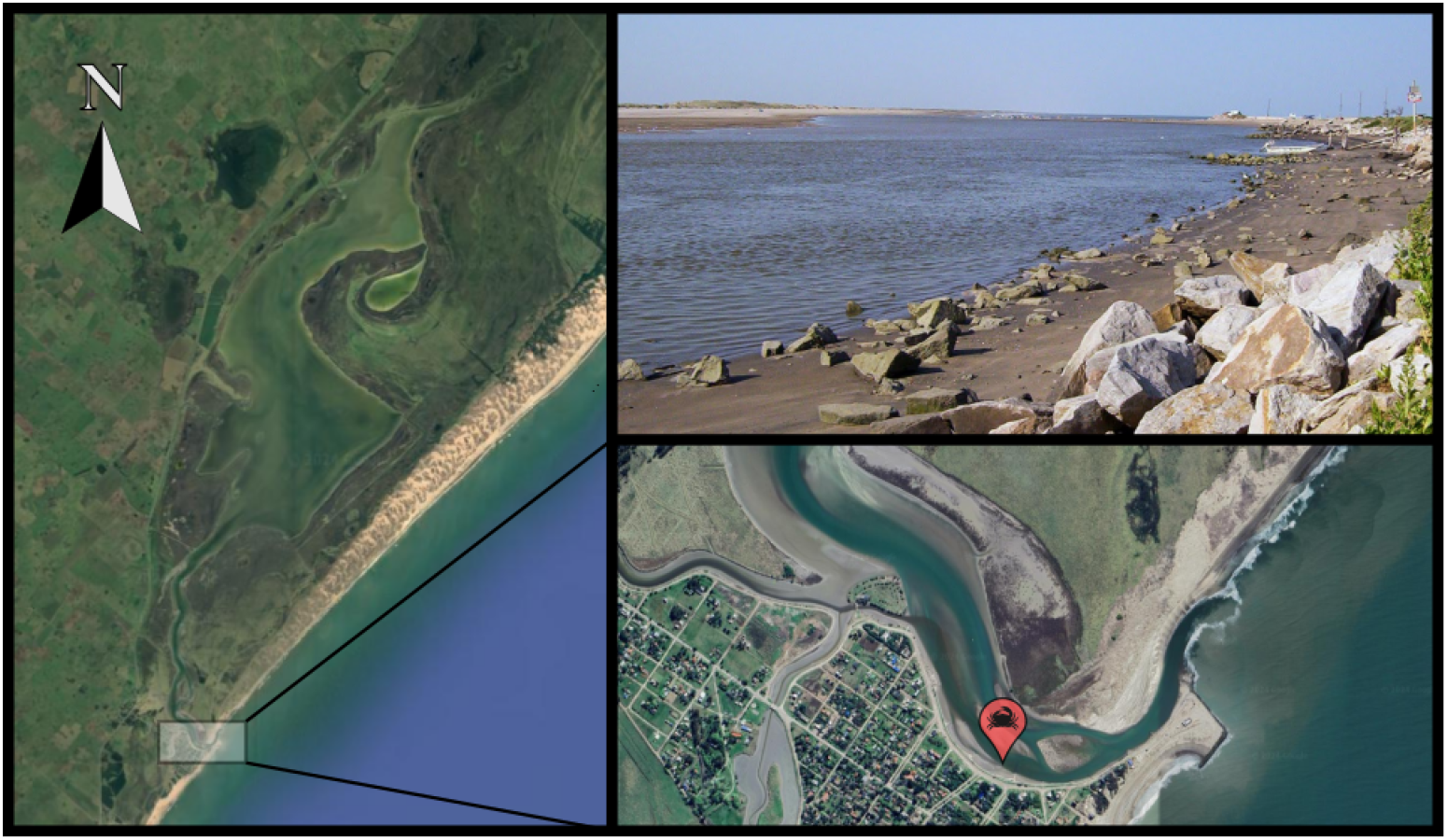
Sampling site at Mar Chiquita coastal lagoon (37° 35’S, 57° 26’W), Buenos Aires, Argentina.

### Sexual differences in body size, relative growth and size at first maturity

Once in laboratory the specimens were sexed, the presence of eggs in females recorded, and the carapace width (CW, hereafter simply referred as “size”), pleon width (PW) and crusher chela length (CrL) measured to the nearest 0.1 mm. Sexual differences in size frequency distributions were tested by applying and comparing Kernel density estimators (KDEs) to the size frequency distributions for each sex (Farias *et al*., 2014). Briefly, KDEs are a non-parametric way to estimate the probability density function of a random variable. This method is more informative since it is sensitive to differences in both the shape of the distribution and its position on the horizontal axis. To test for differences given by the shape of the size distributions only, frequencies were standardized by median and variance (y = x - median/stdev) and then compared. Given the apparent multimodal nature of the size frequency distribution of males upon visual inspection, a Finite Mixed Models analysis was further applied using the MixR package in R (Yu, 2021).

For interspecific comparative purposes the sexual dimorphism was estimated alternatively as

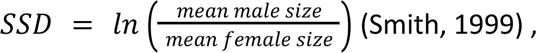

and using the index of Lovich and Gibbons,

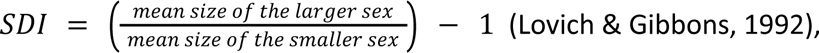

for both the population (*SDI_mp_*) and copulating experimental pairs only (*SDI_mpc_*).

Handedness (namely, in heterochelous species, a bias towards carrying the larger chela at either the right or the left side of the body) was tested using a Chi test with an expected 1:1 ratio for both sexes.

Sexual maturity in both sexes was assessed by the presence of spermatophores in males’ vasa deferentia and the morphology of the females’ abdomen respectively. With the proportion of mature/immature individuals we calculated the size at first maturity as the size at which it is expected that 50% of the individuals are mature (CW_50_).

### Female fecundity and reproductive output

The carapace width of ovigerous females was measured, and the egg masses extracted. The developmental stage of the embryos was determined for 80 eggs from each mass, following previous classifications (Bas and Spivak, 2000; Silva et al., 2003). Fecundity, defined as the total number of eggs per clutch of females with embryos in the initial stage of development, was calculated from the dry weight of the egg mass and the average individual egg dry weight, which was determined using subsamples of 500 eggs from different females. The subsamples were rinsed with distilled water, placed into pre-weighed aluminum cartridges, dried at 80 °C for 48 hours, and then weighed on an analytical balance (with a precision of 0.01 mg). Dry weight measurements were also taken for the whole egg masses and females. Reproductive output (RO) was calculated using the dry weights of the eggs and females, according to established protocols (Clarke et al., 1991; Luppi et al., 1997).

### Mating, male sperm delivery and female sperm storage patterns

Males and females were carried to the laboratory and held separated by sex in 50 l aquaria with 23‰ aerated water at 20 °C and 12:12 (L:D) photoperiod during three days prior to start the experiments. The experiments were conducted in 1-liter cylindrical plastic jars, each containing a mating pair consisting of a mature, non-gravid female and a mature male. Preliminary trials showed that pairs with females larger than males failed to mate. Consequently, pairs were formed with males equally sized or larger than females to ensure mating opportunities, while spanning the entire size combination range to permit the evaluation of potential size-assortative mating. During each trial, six mating pairs were observed simultaneously for 24 hours using a camera connected to a computer. The resulting videos were carefully analyzed to record mating status (whether the pairs mated or not), the time it took for copulation to begin, the duration of copulation, and other behavioral observations. Mating was considered to have occurred when the female was positioned upside down under the male, while copulation was considered to have started when the male’s pleon was enveloped by the female’s abdomen.

At the end of each trial, the reproductive tissue of selected pairs was dissected, regardless of whether copulation had occurred or not. The testes of males were extracted, and the weight of the sperm mass was measured. Females’ ovaries were classified into one of three stages (S1, developing; S2, ready for ovulation; S3, recovering) based on macroscopic features (Ituarte *et al*., 2004). Additionally, one seminal receptacle was extracted and weighed, while the second was examined for the presence of sperm and classified as nsp (no sperm present), sp- (remnants or scarce sperm present), or sp (presence of sperm). The weight measurements are always expressed in milligrams of wet tissue.

To assess the capacity of males and females to deliver and store sperm, we calculated the average maximum deliverable sperm (MDS) and the average sperm mass stored (SMS) for males and females respectively, both for individuals that copulated and those that did not. MDS was determined as the wet weight of the vas deferens, while SMS was calculated as the wet weight of the female seminal receptacle contents. To evaluate if females may store sperm from multiple mates, we applied a t-test to the hypothesis that SMS>MDS in pairs that have copulated.

To assess the patterns of sperm delivery and storage, mature individuals were chosen so as to encompass evenly the size-range observed. The wet weight (WW) of male vasa deferentia, or in females, the combined weight of both seminal receptacles (SR), were recorded using a digital scale with a precision of 0.01 g. Body weight always includes both chelae. For SRs classified as full or partially filled, the structure and contents were weighed separately. To compare the sperm delivery and storage capacities of males and females respectively, the maximum deliverable sperm from a single male (MDS=wet weight of vas deferens) and the total sperm mass stored from an individual female (SMS=wet weight of female SR contents) were determined. The likelihood that the entire volume of sperm stored in the SR comes from multiple mates was assessed using a one-sided two-sample Kolmogorov–Smirnov test of SMS>MDS, under the null hypothesis of identity of the two distributions.

### Overall morphology of the copulatory organs and female seminal receptacles

Among the females collected but not utilized in the experiments, a subset was selected for external and internal morphological examination and analysis of sperm contents. Individuals were classified into different categories based on their abdomen and vulvae morphology and the presence or absence of egg masses. The study of seminal receptacles involved categorizing females into three groups based on their reproductive status: immature females, ovigerous females, and non-ovigerous females.

The complete reproductive system was dissected, observed, and photographed without any additional treatment. Mature females’ seminal receptacles were categorized as empty, intermediate, or full, depending on their size. Additionally, a schematic representation of the female seminal receptacle was created using an Olympus CZX7 stereomicroscope equipped with a camera lucida. To examine the sperm contents, they were delicately extracted by gently opening the receptacle lining and spreading it over a slide. The extracted sperm were then scrutinized under an Olympus CH30 microscope.

## RESULTS

### Sexual differences in body size, relative growth and size at first maturity

A total of 433 crabs (233 males; 200 females) were collected and measured. Once standardized by carapace width, size distribution between sexes differed significantly in position (p<0.001), but not in shape (p=0.344) (Fig. 2). Given the apparent multimodal nature of the size frequency distribution upon visual inspection, a Finite Mixed Models analysis was applied, revealing that the distribution is more likely bimodal (see Supplementary Material S2). Overall, males exhibit a considerably larger size than females (KDE, p<0.001) with *SSD*=0.325, *SDImp*=-0.384 and *SDImpc*=-0.52. Body dimensions measured and the parameters of the allometric functions fitted to each sex are provided in Table 1. Both female and male possess a crushing claw on their right side and a cutting claw on their left side (p<0.0001), although males showed faster relative growth of the crusher chelae. Females’ abdomen grew faster as expected in Brachyura.

**Fig 2.**
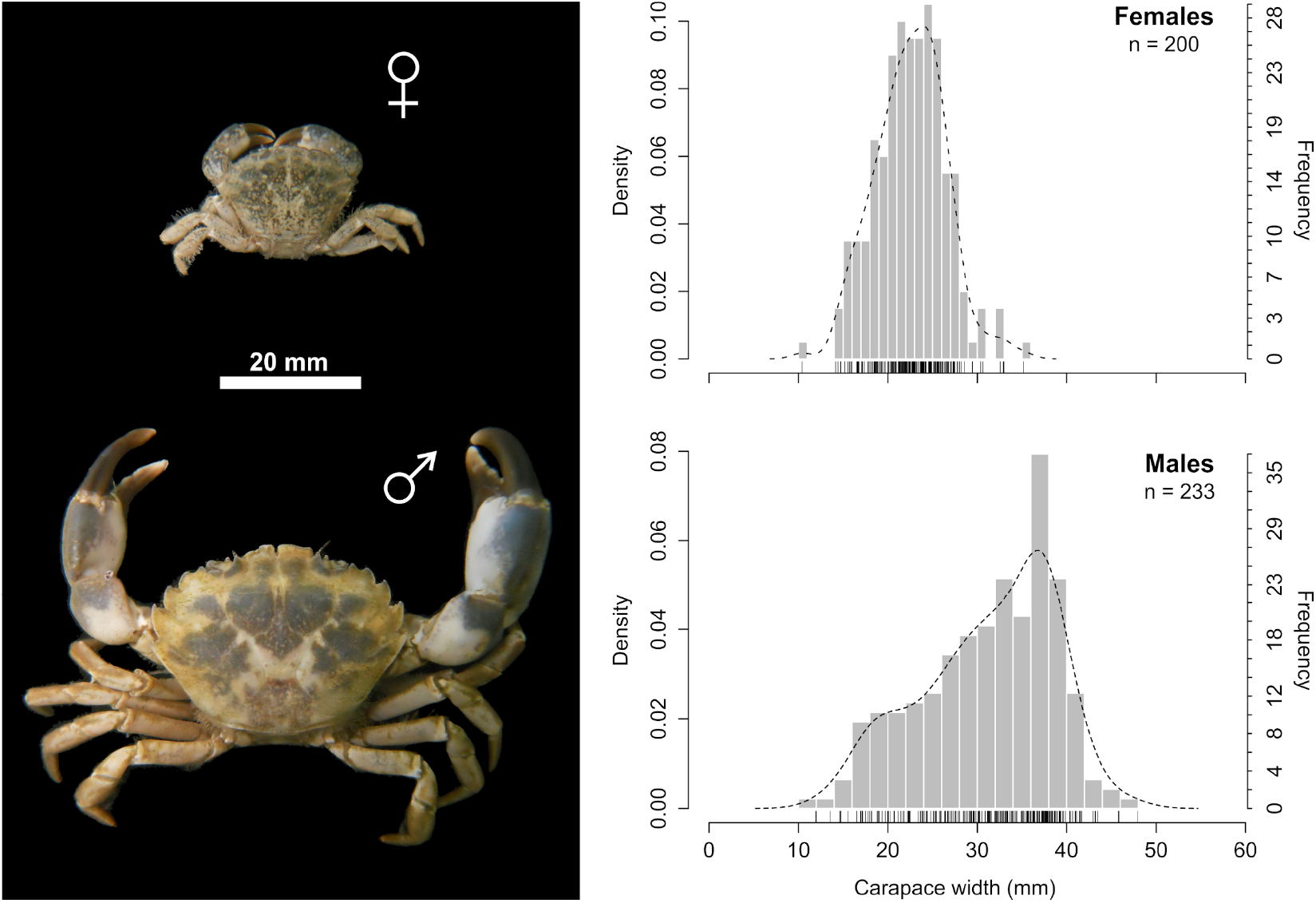
Size distribution of females and males of *Panopeus meridionalis*. Left: Comparative view of average sized male and female in the population sampled. Right: Size frequency distribution of sampled males and females and their respective KDE estimates.

**Table 1:**
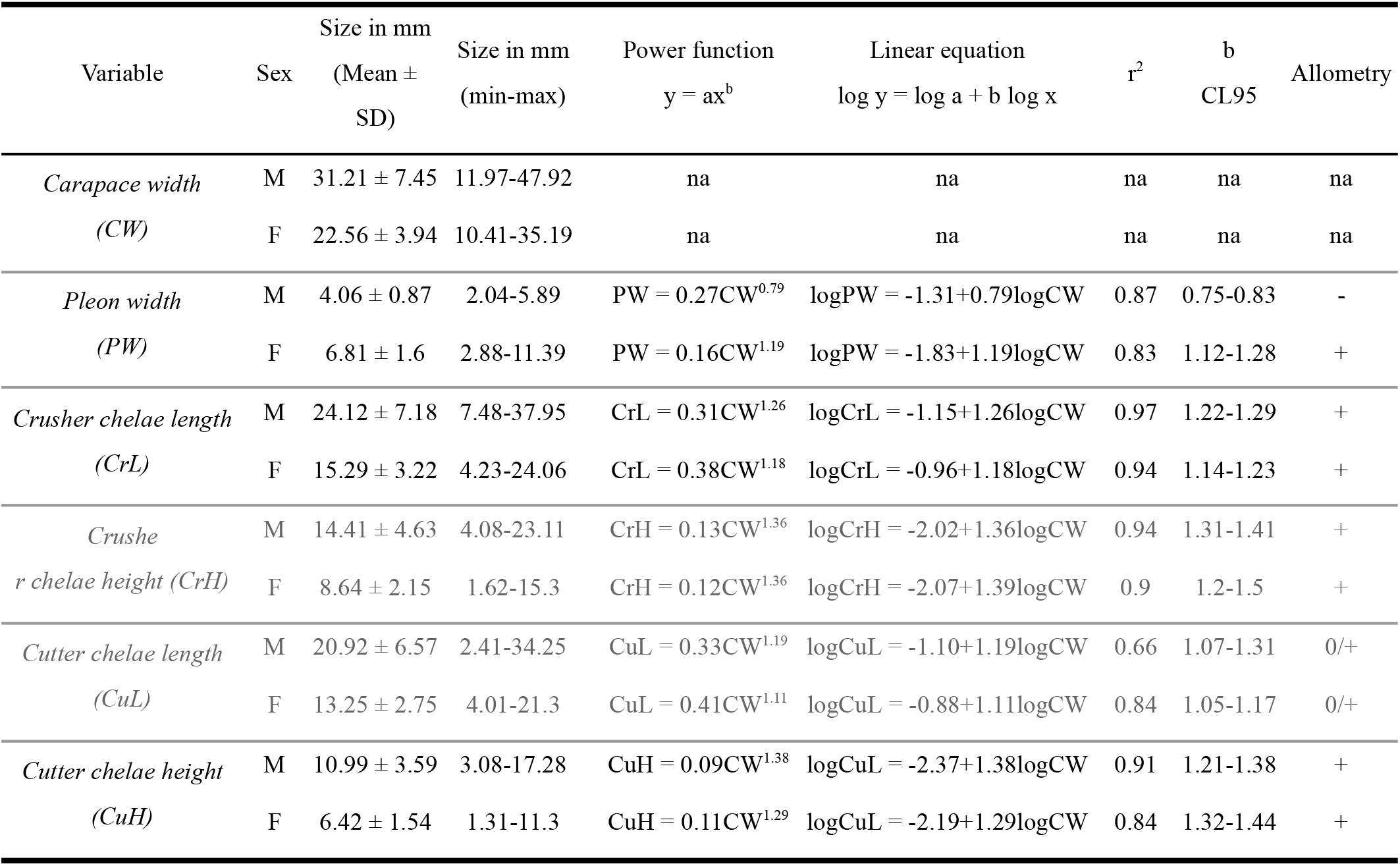
Morphometric variables measured in *Panopeus meridionalis* from Mar Chiquitás coastal lagoon, Argentina, and fitted relative growth functions (as allometry, *sensu stricto*). 95% confidence limits (CL95) are provided for the allometric coeficient “b”.

The size at first sexual maturity was determined to be 21.05 mm carapace width (CW) for males and 19.32 mm CW for females, based on the relative growth of gonopods and the presence of eggs, respectively.

### Female fecundity and reproductive output

To assess fecundity and reproductive output, 24 ovigerous females with carapace width ranging from 19.32 to 32.9 mm were examined. The number of eggs was found to be proportional to carapace width (Fig. 3a). The average fecundity of the females (mean±SD) was 28925±16276 eggs/female, and the average reproductive output was 0.081±0.022 (8.1%). Notably, the reproductive output of *P. meridionalis* was independent of crab’s size (Fig. 3b).

**Fig 3.**
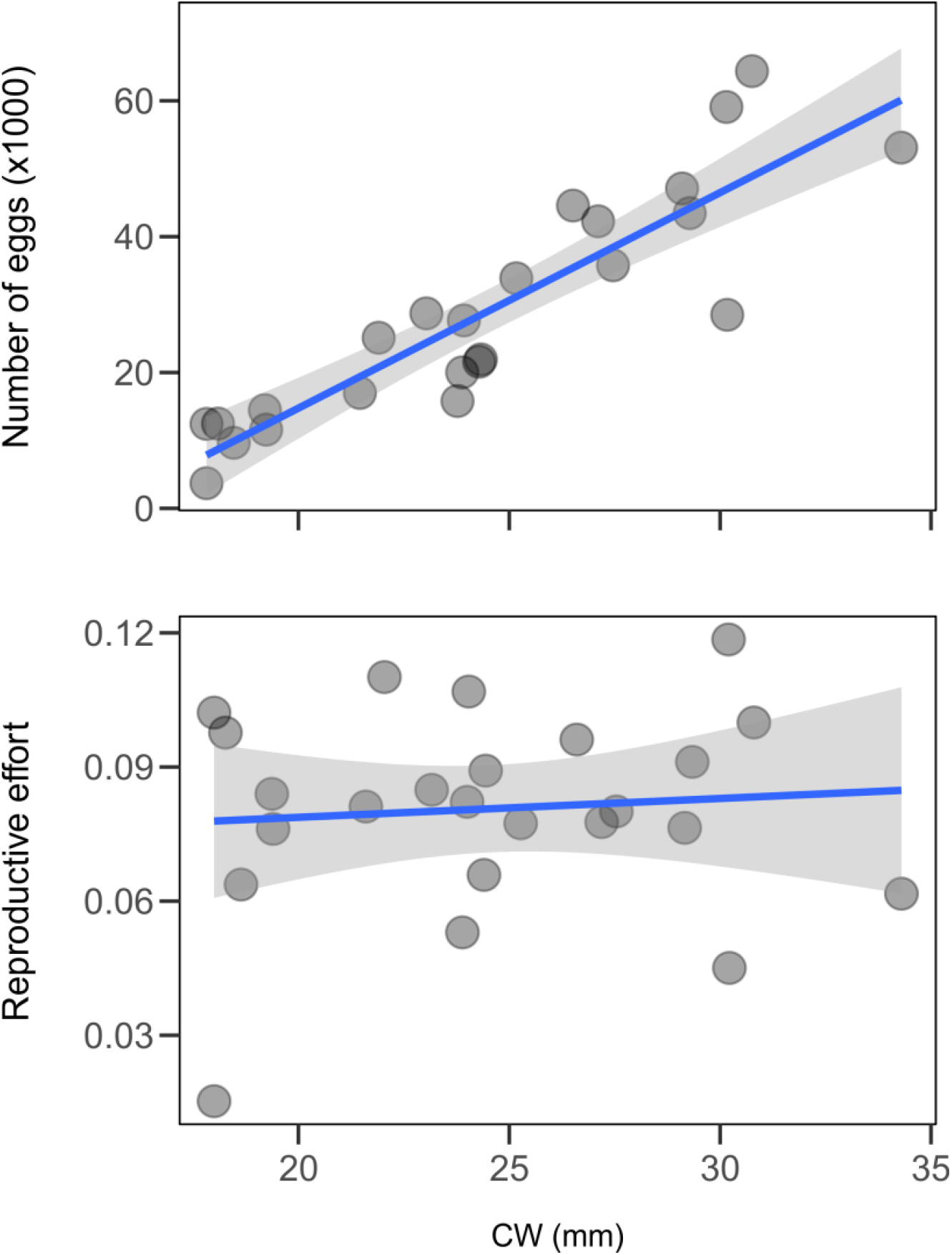
Regressions for the relationship between carapace width and fecundity (top), and reproductive output (bottom), of female *Panopeus meridionalis*.

### Mating, male sperm delivery and female sperm storage patterns

A total of 117 experimental pairings were conducted, resulting in copulation in 19 instances (∼16%). In all cases, copulation occurred in pairs where the male was conspicuously larger, and females were hard-shelled (Fig. 4a). Differences in SSD between pairs that copulate and those who did not were tested by a one-sided Bootstrap Welch Two Sample t-test that yielded a *p*-value of 0.11, indicating that pairs that engaged in copulation had males larger than their female partner. The smallest SSD recorded in successfully copulating pairs was 1.3, which says that the male had a carapace width 30% larger than its partner. Coincidently a two-sided Welch Two Sample t-test for differences in the slope and intercepts of simple regression models applied to the size ratio of the pairs in both groups (Fig. 4b), resulted in equal slopes (two-sided t-test, *p*=0.08) but higher intercept of pairs that copulated (one-sided t-test, *p*=0.02).

**Fig 4.**
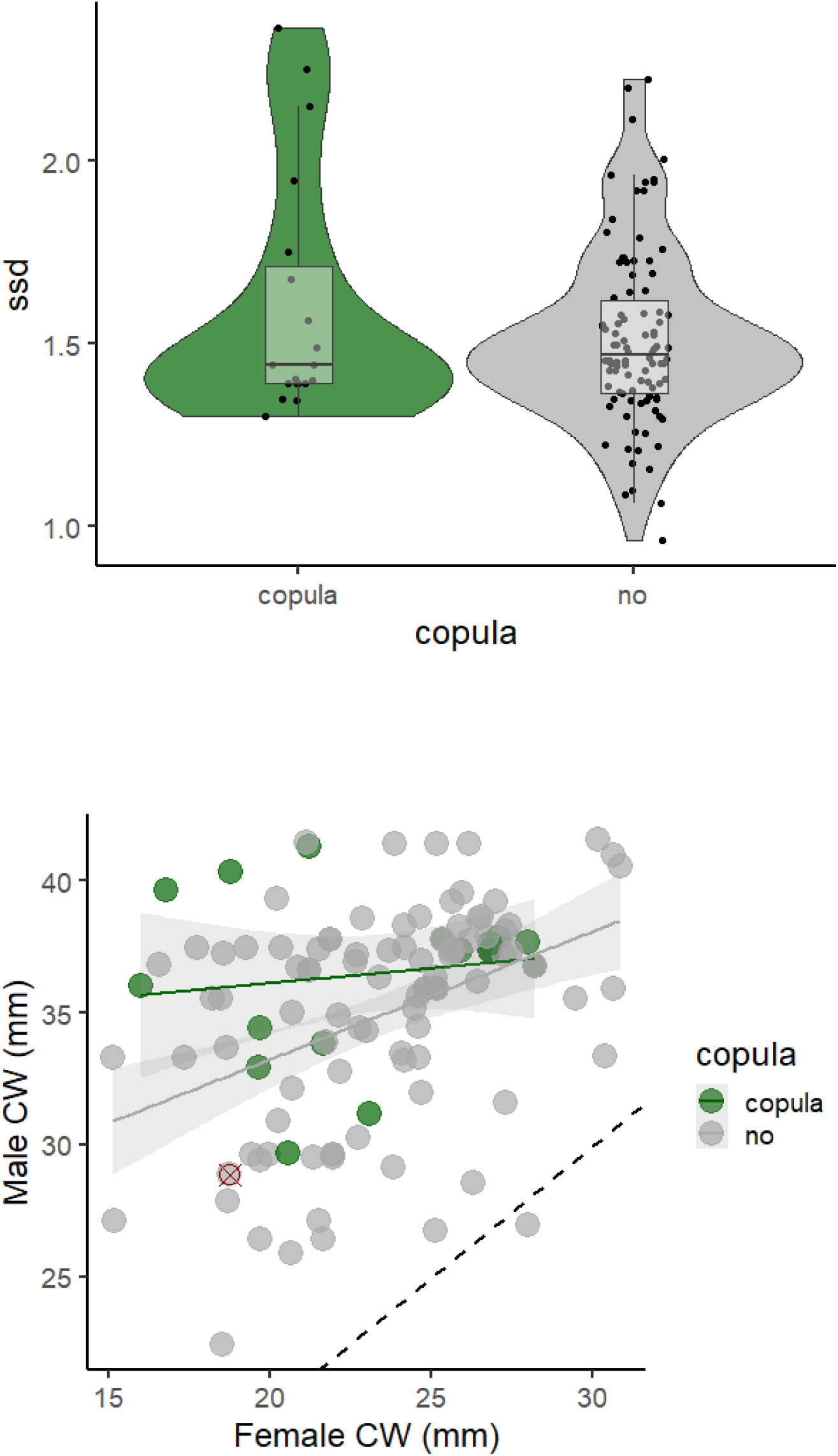
**a**) Distribution of sexual size dimorphism (SSD) for experimental pairs that copulated and those who didńt. Pairs with SSD below 1.3 did not copulate. **b**) Size relation between males and females of *Panopeus meridionalis* used in mating experiments. N pairs=117. Grey dots: pairs that did not copulate, Green dots: pairs that copulated. red cross = Experimental pair where the male ate its female partner. Dashed line: male size equals female size.

The majority of females analyzed possessed ovaries in stage 1 (36%) or stage 2 (44%) (Fig. 5a), of whom 86% had seminal receptacles containing varying amounts of sperm (Fig. 5b). Among females that copulated in the lab and those who did not, 89% and 76.8% had ovaries in stage 1 or 2 respectively. In both groups, the majority of females had spermatozoa or detectable remains of sperm in their seminal receptacles (88,2% in females that copulated and 85.8% in those who did not). The probability of an experimental pair engaging in copulation was independent of both the ovary stage and the presence of sperm in the female seminal receptacles (Fisher’s exact test of independence with *p*-values of 0.53 and 0.66, respectively).

**Fig 5.**
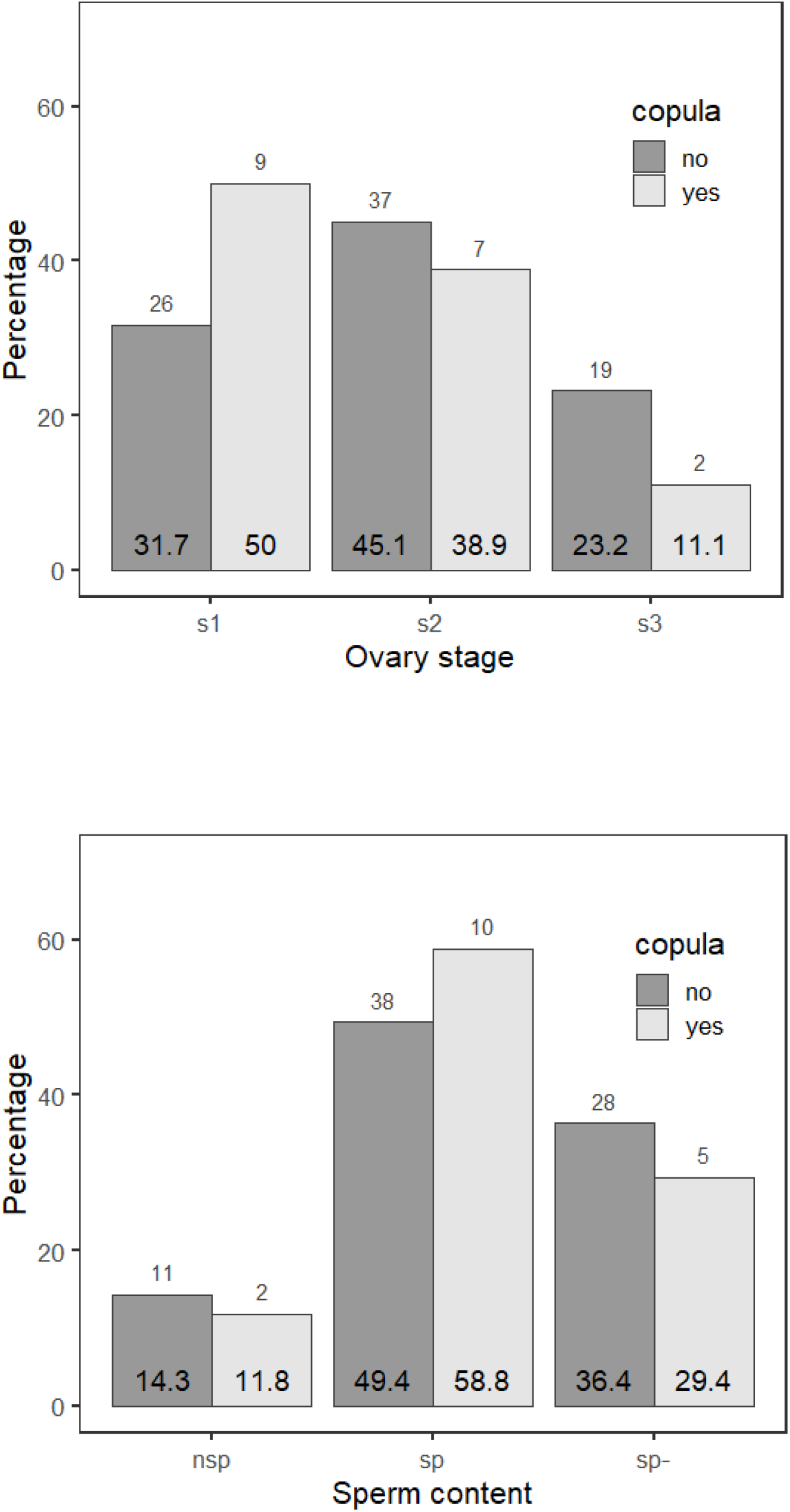
Percent of *Panopeus meridionalis* females of **a**) given ovary stage (1 = developing; 2 = ready for ovulation and 3 = recovering) and **b**) sperm contents in seminal receptacles (nsp = no sperm; sp- = scarce sperm and sp = presence of sperm) for mated and unmated crabs. Numbers above the bars are individuals in each category. The numbers inside the bars are the calculated percentages of number of individuals in the category relative to the total in the same subcategory (bars of the same color in each plot)

The average time of pre-copulatory activities (time from the first attempt of grabbing the female until copulation) was 18 minutes, usually involving some failed attempts, final capture, and positioning of the female for gonopod intromission. The male exhibited a distinctly coercive mating behavior during this period. At times, the male slowly approached the female, grabbing her chela or leg with a fast movement when she attempted to escape. Alternatively, the male sat-and-waited and rapidly captured the female when she passed nearby. In most instances, females react by fully extending both chelae in the typical defensive posture of crabs. When males persist, the females become motionless while the male attempts to manipulate and position her upside-down beneath its body for copulation (supplementary material S1).

In all instances, crabs kept the ventral-facing position with the male placed over the female. Typically, the female appears to resist briefly but repeatedly with rapid movements while positioned beneath the male. Eventually the female stops resisting and retracts her legs and chelae. Rhythmic movements of the male pleon were observed during copulation. The end of copulation was marked by an abrupt release of the female with no evidence for subsequent post-copulatory mate guarding. If copulation lasts, females often attempt to escape, with varying success. Some females succeeded to break free, but in most cases the male managed to relocate the female back to the original mating position. The duration of copulation, including interruptions, varied between 11 and 381 minutes, with an average of 194 minutes, and was not correlated with the degree of size dimorphism of the mating pair (p=0.17; Fig. 6).

**Fig 6.**
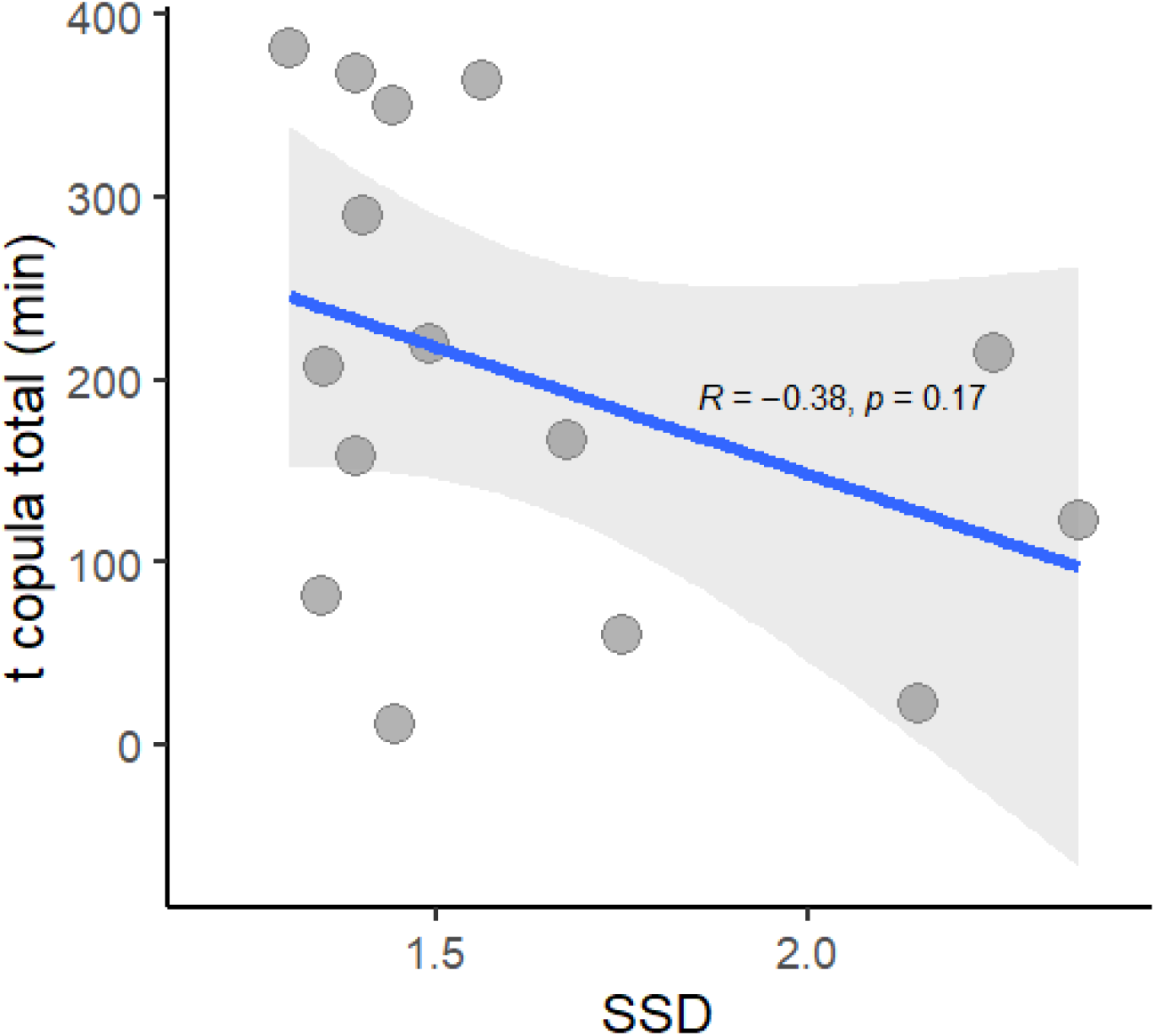
Simple regression for the relationship between sexual size dimorphism (SSD) of the mating pair and duration of copula in *Panopeus meridionalis*.

The maximum deliverable sperm (MDS) of males that did not copulate was 0.26±0.16 mg, while for copulating males, it was 0.36±0.11 mg. Copulating males exhibited a 27.7% higher sperm content than non-copulating males. The SMS for females that did not copulate was 0.012±0.01 mg, whereas for copulating females, it was 0.042±0.028 mg. Females engaged in copulation exhibited a 71.4% greater sperm mass storage (SMS) in their seminal receptacles compared to non-copulating females, as expected. The hypothesis that mean MDS = mean SMS was rejected (t-test, p<0.001). The MDS/SMS ratio between copulating males and females was 8.53.

### Structure of the female reproductive tract

The female reproductive tract of *P. meridionalis* follows the typical brachyuran pattern, comprising a paired structure, each consisting of a vulva, a vagina, a seminal receptacle for sperm storage, and an oviduct connecting the latter with the ovary. The seminal receptacle is a spherical sac characterized by a dorsal, glandular, flexible portion and a ventral chitinous base. The vagina (Va in Fig. 7) is a straight tubular structure with a basal portion featuring a wall that is partly thick and rigid and partly thin and flexible. The vagina extends dorsoventrally, from the seminal receptacle to the thorax, where it forms a flexible fold, the vulva (Vu in Fig. 7), which closes the passage to the exterior. The terminal portion of the ovary (labeled as ‘Ov’ in Fig. 7) connects to the dorsal tip of the seminal receptacle. No spermatophores were found within the seminal receptacles, either in experimentally mated females or in those captured in the field.

**Fig 7.**
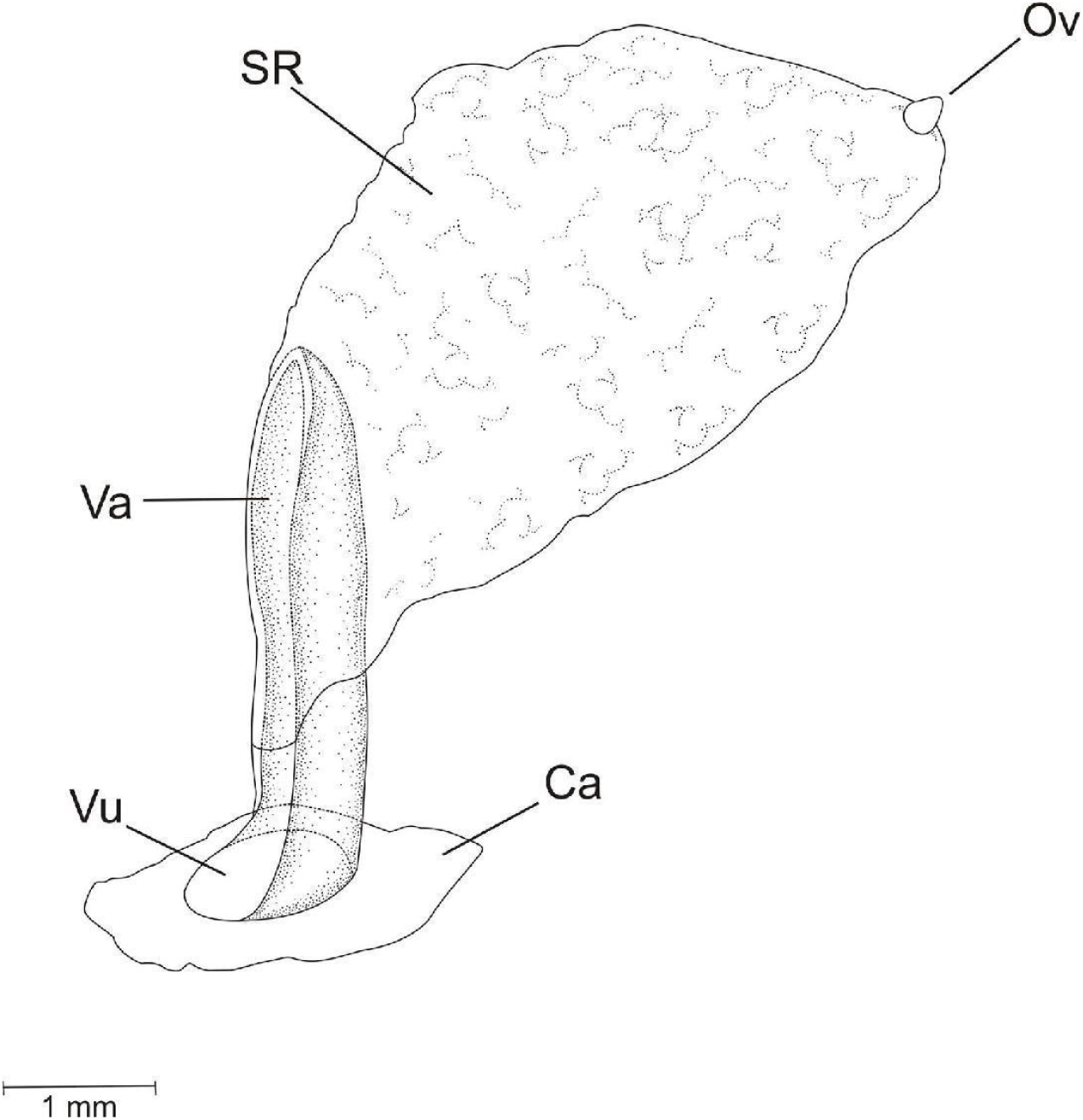
*Panopeus meridionalis* seminal receptacle structure. Ov: dorsal ovary connection, SR: seminal receptacle, Va: vagina; Vu: vulva; Ca: ventral part of the carapace (6th sternite)

## DISCUSSION

### Sexual differences in body size, relative growth and size at first maturity

The male biased sexual size dimorphism (MBSSD) of *P. meridionalis* ranks among the highest reported in animals, jointly with that of other crustaceans, some primates, mustelids and ungulates (Fairbairn, 1997; Janicke & Fromonteil, 2021). While the evolution of larger males is often attributed to enhanced performance in male-male competition for mating and female preference, the fact that the larger males are better able to force females to copulate is often overlooked as a concomitant powerful evolutinary driver (Shine & Mason, 2005; Wilhelm *et al*., 2015). Although we did not test for the size related outcome of male-male contests, there is a great deal of evidence that positive correlation between body size and hierarchy is almost ubiquitous among crabs (Swartz, 1976; Van Der Meeren, 1994; Sneddon *et al*., 1997; Brown *et al*., 2005; Daleo *et al*., 2009; Sal Moyano *et al*., 2016) so it is reasonable to think that *P. meridionalis* is not an exception and intrasexual competence is operating. However, given the large size difference and aggressive mating behavior of males, it is likely that overt sexually antagonistic coevolution (Eberhard, 2004) play a major role in driving the sexual dimorphism in this species, although in reciprocal reinforcement with sexually antagonistic selection (Connallon & Clark, 2014). In tandem with body size, differences in relative growth show rapid evolution of male weaponry, which in crabs is also involved in increasing male’s opportunities to access females and enhanced intrasexual competition (Shuster & Wade, 2003; Baeza & Thiel, 2007; Baeza & Asorey, 2012).

Contrary to the expected for natural populations with standard dynamics, where smaller (younger) individuals are in larger numbers than larger (older) individuals, the smaller males were conspicuously underrepresented (see figure 2). This does not appear to result from sampling bias, as the females of the same size range of the smaller males were captured in roughly equal numbers to those of the males in general. Instead, it suggests either the accumulation of females in the narrower size range determined by the lower growth rate and maximum size compared to males, or the segregation of smaller males to habitats or low tidal levels beyond the scope of our sampling. Considering the aggressiveness observed in larger males during the experimental pairings, and the described in other crab species from similar environments (Allen *et al*., 2010; Casariego *et al*., 2011), the displacement of smaller males to peripheral areas to avoid potential harm from male-male antagonisms is plausible. Male harassment of females may also promote sexual segregation in habitat use, but since male-female interactions are typically less intense than intrasexual antagonisms, it also can lead females to coexistence with larger males, particularly during the reproductive period (Croft *et al*., 2006; Darden & Croft, 2008).

Lastly, it is noteworthy that the size range at which males become sexually mature overlaps with the average size of the co-occurring females. That is, most males attain sexual maturity at a size that enhances their chances to catch and manipulate females for mating, as observed in the experimental pairings. This fact reinforces the idea of spatial segregation of males based on their size (and maturity stage). In this context, we anticipate that females co-occurring in the field with larger males would be mature and receptive, which usually mitigates male aggression upon encounter and likely moderate other indirect short and long-term adverse effects of cohabitation in the context of sexual coercion (Makowicz & Schlupp, 2013; Iglesias-Carrasco *et al*., 2019). This holds especially true if this spatial coexistence is limited to the reproductive period, though it remains a prediction yet to be tested.

### Female fecundity and reproductive output

In females, a great deal of the variability in parental investment lies in the number of eggs produced. Males tend to fertilize most or all of the eggs laid by the female, but there is no guarantee that this will contribute to succeeding clutches. This results in sexual conflict over clutch size: while maximizing the number of eggs laid soon after copulation enhances the male’s fitness, the female may optimize her overall lifetime reproductive output by restricting the size of the current clutch, prioritizing subsequent survival and, in turn, influencing the number of future clutches. Figure 3a shows that in *P. meridionalis*, brood weight and the number of eggs per brood are positively correlated with female body size, which aligns with the fecundity advantage hypothesis that predicts selection towards larger females because they lay more eggs. However, the pronounced MBSSD of *P. meridionalis*, suggests that the selection for larger, more fecund females is heavily counteracted by opposing forces, likely the high intraspecific aggression that is well-known to exacerbate when the antagonists are more similar in size. In contrast to fecundity, the reproductive effort (the fraction of body mass devoted to reproduction) appears constrained to approximately one tenth of female body weight, in agreement with what was described as the rule for free living true crabs given the limited space available for yolk accumulation within the female cephalothorax (Hines, 1992). Hence, considering the absence of correlation between reproductive effort and female size (Fig. 3b), it appears that the overall reproductive benefits of being larger do not outweigh the associated costs of heightened male harassment and derived consequences like the displacement of females to lower-quality areas or reduced feeding rates (Pilastro *et al*., 2003), among others.

### Mating, male sperm delivery and female sperm storage patterns

Experimental pairings revealed that mating in *P. meridionalis* involves strong male harassment, possibly including forced copulation (see the next section and video in supplementary material S1) with no apparent pre or post copulatory guarding. This observed mating behavior closely mirrors that described by Swartz (1976) for *Dyspanopeus sayi* (Smith, 1869). In experimental pairs, all females, including those that mated successfully (’receptive females’), actively avoided the male and, when captured, resisted vigorously. In turn, males exhibited active pursuit of the female or employed a sit-and-wait strategy, capturing her with sudden claw movements, similar with the described in *P. herbstii* for catching mobile prey (Silliman et al., 2004). Our simple experimental setup eliminated male-male interferences and, due to confinement, significantly reduced female escape opportunities. Despite these conditions, many males attempting pursuit failed to capture, position and mate with the females. According to Clutton-Brock and Parker (1995), females may enhance their fitness by delaying or refusing mating, enabling pairing with a superior male or breeding under more favorable ecological circumstances. We anticipate that femalés avoidance of males would constitute an important aspect of the mating system of *P. meridionalis* to take into account when studying size and sex spatial structure in nature.

Except for Swartz’s (1976), prior studies on other panopeids, *Eurypanopeus depressus* (Smith, 1869) and *Rhithropanopeus harrisii* (Gould, 1841), reported none or inconspicuous pre-copulatory behaviors (Rodgers *et al*., 2011). In our study, successful male capture induced a drastic shift in female behavior from avoiding/resistance to submission. These phases may represent distinct processes of a rudimentary form of precopulatory behavior, where female resistance functions as a mechanism for cryptic mate choice (Eberhard, 1996, p. 199), enabling the evaluation of male quality through its ability to coerce (Chapman et al., 2003). Alternatively, the female’s sudden immobility might represent a ‘playing dead’ behavior, often observed in response to predators, but in this context, triggered by male harassment during mating. After the male overpowers the female’s resistance, the playing dead response likely serves as a ritualized strategy to prevent injuries or potential harm, a common outcome in coercive mating systems (Chapman *et al*., 2003; Issa & Edwards, 2006). Accordingly, in our experiments successful mating occurred only with those males large enough to subdue the female (males more than 30% larger than the female). Large males have the potential to inflict serious injuries to females. During the experimental matings we have recorded cases in which the male crushed and killed the female, and one case in which the male eventually ate his partner (red dot in Fig. 4b). Inflicting costs onto females that do not cooperate is a widespread phenomenon that may increase male mating success in coercive mating systems (Clutton-Brock & Parker, 1995), and might be operating here.

Even in the scenario of overt sexual coercion, in the long run the females may benefit by passing increased survival, mating success, and fitness associated with dominant males to offspring, (the sexy sons or the good genes hypotheses) so that a submission can also be interpreted as part of the repertoire of behaviors available for female choice in this context (Snow *et al*., 2019; Snow & Prum, 2023). Accordingly, the fact that the experimentally mated females all had developing or ready to spawn ovaries (Fig. 5a) suggests that females are more receptive (or resist less) when in this stage, strengthening the idea that the submissive behavior described here may be a way to gain some control over the reproductive outcome. An alternative but not mutually exclusive explanation is that there is some mechanism by which males can sense female receptivity and hence invert more in subduing the female. There is evidence suggesting that even under coercion, females may retain the opportunity to indirectly choose mates through the release of sex pheromones (Okamura & Goshima, 2010), although that is frequently accompanied by other morpho-physiological changes related to receptivity, such as the softening of the vulvae opercula (Sal Moyano *et al*., 2017), a trait not observed in these species. Nevertheless, given the turbidity of the shallow waters inhabited by *P. meridionalis* chemical cues probably play a role in intraspecies agonistic interactions and further study is required on this (Rodgers *et al*., 2011).

A last interesting fact requiring interpretation was the great differences in time invested in copula that we registered during the experimental pairing (Fig. 6). It might be related to the mating history of the male, the female, or both, but also derived from individual differences in the level of female resistance and male endurance, as demonstrated experimentally in some insects (Arnqvist & Rowe, 1995; Jormalainen *et al*., 2001). However, it is important to keep in mind that in nature, the mean interaction time may be much shorter given that experimental conditions of confinement prevent the female from escaping when breaks away from her partner.

Contrary to inferred for *P. herbstii* based in individual and pair distribution (Silliman *et al*., 2004), mating in *P. meridionalis* certainly involves hard females. We cannot rule out mating with soft females, as occurs in many other brachyuran crabs, but it is hardly a chance for the female given the aggressive behavior of males during mating. The hard shelled mating and the lack of pre and post copulatory mate guarding along with the relatively short copulation time, suggest that post-mating sexual selection, mainly sperm competition, does not play a significant role (if any) in the mating system of *P. meridionalis*.

### Structure of the female reproductive tract

In scenarios with the potential for forced insemination or limited mate selection, sexual conflict often extends to postcopulatory mechanisms. Females are predicted to evolve anatomical and physiological adaptations to manipulate fertilization (cryptic female choice), thereby countering fitness costs from male coercion (Stockley, 1997; Eberhard, 2009; Parker *et al*., 2010). Brachyuran crabs have evolved a striking diversity of female morpho-physiological adaptations to control insemination and the fate of male ejaculates in response to the different species-specific contexts (Diesel, 1991; McLay & López Greco, 2011; McLay & Becker, 2015). Different species have evolved physical barriers to prevent forced insemination, such as movable vulvar opercula attached to muscles that permit occluding the reproductive ducts at will, or external rigid lids or plugs that become flexible during the reproductive period. Females also evolved seminal receptacles with diverse morpho-physiological adaptations that may function in long-term sperm storing, expulsion of sperm masses after mating, denature or digest spermatozoa, encapsulate it, and even manipulate the order in which different sperm masses from sequential matings encounter and fertilize the eggs (Sal Moyano *et al*., 2010; Becker *et al*., 2011; Pardo *et al*., 2013; Klaus *et al*., 2014; Vehof *et al*., 2014; Becker & Scholtz, 2016; Farias *et al*., 2017). On its side, males evolved gonopods with varied species specific morphological features (Becker *et al*., 2012) likely aimed to prevent sperm competition and the intensity of polygyny (Pretterebner *et al*., 2022). They also have the ability to deposit their spermatozoa encapsulated in spermatophores that provide nutrition and protection to the cells, allocate the volume of ejaculates based on the socio-sexual context, and even sequentially ejaculate sperm and seminal fluid that hardens within the seminal receptacle in order to isolate or displace sperm masses previously deposited by competing males (Sainte-Marie, 2007; Pardo *et al*., 2016, 2018). The patterns of male sperm delivery and female storage of *P. meridionalis*, as well the structure of the female reproductive tract, resembles partially that described in *D. sayi* (Swartz, 1976) and provides further support for the hypothesis of a coercive mating system and strong sexual antagonistic coevolution at play.

If not duly contextualized, pre-intromission female resistance (as obsereved in our experimental pairings) can be straightforwardly perceived as a behavior to prevent forced copulation, or interpreted as a way of testing male endurance or quality (Painting *et al*., 2016). In insects for instance, male persistence in attempting to force female genitalia open functions as a screening mechanism for suitable mates and is more aptly described as ‘persuasion’ (Eberhard, 2002). However, in *P. meridionalis*, females lack temporarily hardened or movable opercula, or other closing structures that may function to mechanically hinder gonopod penetration as described in other species (Sal Moyano *et al*., 2012; Farias *et al*., 2017). Even considering the female’s resistance as part of a cryptic mate selection mechanism, her chances to repel unwanted copulations seems confined to brief counter aggressions or escaping given that once the female is grabbed by the larger male, has little chance to resist without being harmed. Thus, lacking evidence to the contrary, the female behavioral responses to male attempts in *P. meridionalis* is more accurately described as female resistance to coercion rather than any alternative emphasizing female cooperation.

Under coercive dynamics it may be expected a small capacity for female sperm storage because mating typically involves brief, forced copulations, limiting the opportunity for females to choose mates and to store sperm over an extended period. As a result, female mating tactics and anatomical adaptations are expeted to be geared towards efficient and rapid sperm usage. Indeed, the seminal receptacles of *P. meridionalis* appear as a pair of fairly simple sacs (Fig. 7), lacking the capability of elongation and the more complex structures associated with sperm long-term storage, manipulation and postcopulatory selection observed in several other brachyurans (Diesel, 1991; Farias et al., 2017; McLay and Becker, 2015). Moreover, the dorsal connection between the ovary and seminal receptacle, positioned opposite to the entry of the vagina (Fig. 7), indicates enhanced fertilization opportunities for the initial (or the only) male to mate with the female if sperm masses stacks consecutively (Diesel, 1991; McLay & López Greco, 2011a); or alternatively, if sperm mixes within the receptacles, it implies equal chances of fertilization.

In both scenarios, males increase their fertilization chances by being the first to mate with as many females as possible within the shortest time and by filling up the receptacles to exclude sperm from other males. The observed absence of pre or post-mating guarding behavior aligns with this prediction, as guarding females in this context wouldn’t contribute to securing paternity but rather reduces the time spent searching for new mates. It further aligns with the calculation that average males could fertilize up to eight females before depleting their sperm reserves (*MDS/SDS =* 8), indicating they must be capable of filling up the seminal receptacles and ensuring paternity of several mates if given the opportunity. These findings are also coherent with the expectations of a coercive system where females, lacking direct control over mate selection or specific paternity, would increase mate diversity and reduce costs of coercion by storing the minimal sperm required for each spawn and then remating for the next reproductive cycle. This would be a potent evolutionary force towards smaller body size, counteracting the traditionally invoked fecundity selection for larger females, and accounting for the observed low and simple sperm storage capacity in females. Lastly but not less significant, our study reveals that males transfer free sperm rather than spermatophores, as observed in *Macrophthalmus (Hemiplax) hirtipes* (Jennings *et al*., 2000) and *D. say*i (Swartz, 1976). The absence of spermatophores in the ejaculates has been linked to ephemeral storage by the female, and jointly with the relatively small size of the seminal receptacle (SR) in relation to body size compared to other crab species, further supports the notion that long-term sperm storage is not a feature of the species’ reproductive system.

### Concluding remarks

Here, we present novel information on the reproductive and life history traits of *P. meridionalis* and integrated this information to evaluate our hypothesis that the sexual dimorphism with males being conspicuously larger than females results mainly from antagonistic selection forces driven by sexual conflict, rather than differing natural selection forces between the sexes. The species is characterized by a highly male-biased dimorphism in size and weaponry, no correlation between female size and reproductive effort (the fraction of body mass devoted to reproduction), a (likely ritualized) submissive mating behavior of the female, no pre or post copulatory guarding, small female seminal receptacles with reduced sperm storage capacity and sperm contents consisting mostly on free spermatozoa, lack of morpho-anatomical features that would suggest the existence of post mating selection processes (sperm competition and cryptic female choice), and lastly, absence of anatomical structures of the female that could be interpreted as barriers or deterrents to forced penetration by the male. On the other hand, there is no evidence to suggest that males and females are subject to differential selection forces that are not, in turn, correlated with sexual differences in body size (such as sex differences in predation risk, feeding rates and access to prey). Considering the spatial homogeneity of the estuarine areas shared by both sexes, there are no evident factors of sexually antagonistic selection forces (natural selection acting differentially on each sex) that can be accounted as main drivers for the pronounced dimorphism observed. All together, this constitutes a coherent body of evidence that strongly supports the existence of an intrinsically coercive mating system in the species. However, there is a wealth of information yet to be known to comprehensively grasp the evolutionary processes resulting in a functional coercive system as resolution of sexual conflict. For this species, it includes the study of the connection between mating and molting, sex differences in habitat use, feeding, growth, mating, and mortality rates, the femalés receptivity period, the mechanisms of mate attraction and detection, and quantifying costs of male-male aggression and the fitness effects of male harassment on females in the field, among others. Many of these studies require experimental approaches for thorough investigation.

All together, *Panopeus meridionalis* proves to be an interesting model for further studying the resolution of sexual conflict through antagonistic selection and coevolution taken to extremes, and opens up a promising avenue for research on the relationship between the resolution of evolutionary conflict via a sexually divergent path and the stability and low complexity of the environment in which the species had evolved. The last would be worthwhile to explore further by expanding the study to other panopeid mud crabs, considering the taxon’s morphological stasis and the stability of the mudflats they are adapted and constrained to live in (Schubart *et al*., 2000).

## Acknowledgements

The field work and sampling was partially supported by funds of Consejo Nacional de Investigaciones Científicas y Técnicas (CONICET) (PIP 112-200801-00176) and Universidad Nacional de Mar del Plata (UNMdP: EXA517 /10). NEF would like to dedicate this work to Dr. Christoph Schubart, a dear friend and valued colleague, whose boundless curiosity and kindness inspired many of us to take an interest in the fascinating world of diversity and evolution.

## Supplementary material

**S1:** Result of the test for sexual differences in size frequency distributions (SFD) using Kernel Density Estimators, and exploration of multimodal SFD in males using Finite Mixture Models (Yu Y, 2021)

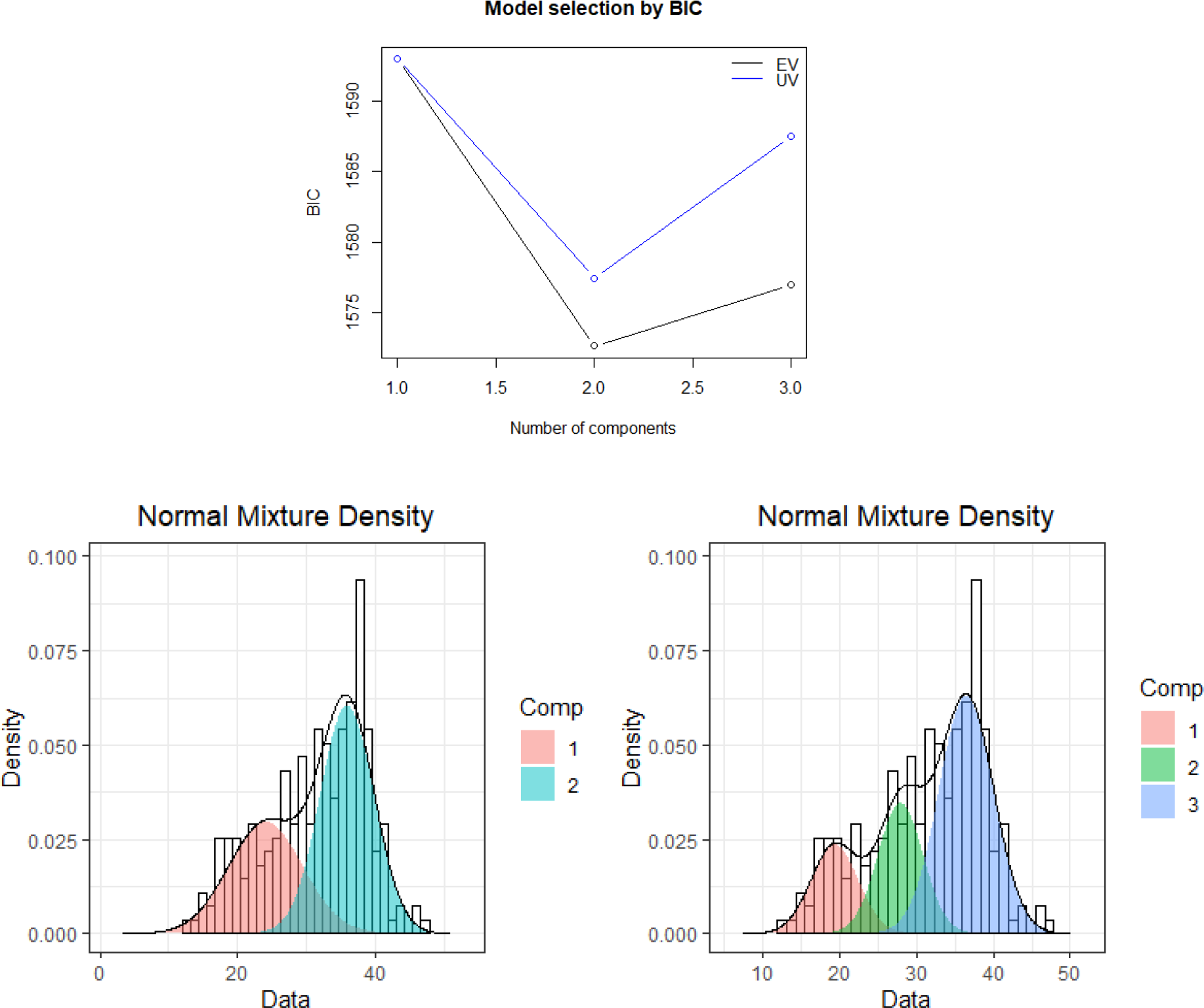

Yu Y (2021). _mixR: Finite Mixture Modeling for Raw and Binned Data_. R package version 0.2.0, <https://CRAN.R-project.org/package=mixR>.

**S2: Mating experiment videos**

<u>120307_2324.AVI</u>

<u>120307_2219.AVI</u>

120315_2341.AVI

